# Tastes and retronasal odours evoke a shared flavour-specific neural code in the human insula

**DOI:** 10.1101/2025.01.06.631354

**Authors:** Putu Agus Khorisantono, Maria G. Veldhuizen, Janina Seubert

**Affiliations:** Division of Psychology, Department of Clinical Neuroscience, Karolinska Institutet, SE-171 77 Stockholm, Sweden; Department of Psychology, Faculty of Science and Letters, Mersin University, Mersin, Türkiye; Department of Anatomy, Faculty of Medicine, Mersin University, Mersin, Türkiye; National Magnetic Resonance Research Center (UMRAM), Bilkent University, Ankara, Türkiye

## Abstract

During food consumption, tastes combine with retronasal odours to form flavour, which leads to a link so robust that retronasal odours can elicit taste sensations without concurrent taste stimulation. However, the cortical integration of these parallel sensory signals remains unclear. Here, we combine a flavour-binding paradigm and functional neuroimaging to test whether retronasal odorants evoke encoding patterns in the insula similar to those of their paired tastants. Healthy participants attended a familiarisation session with congruent sweet and savoury flavours followed by two fMRI sessions where they separately received the constituent tastants and odorants. Multivariate pattern analysis revealed classification of retronasal odours within the insula, exhibiting overlapping representations with their associated tastes, particularly in the ventral anterior insula. Additionally, we observed temporal instability in insular taste representations, paralleling findings in rodent gustatory cortex. These findings underscore the robust integration of gustatory and retronasal olfactory processing that underpin the flavour percept.

## INTRODUCTION

Most of us who have suffered from a cold can attest to the illusory feeling of taste loss that comes with a blocked nose. This happens because normally, when food is in the mouth without a blocked nose, odorants reach the olfactory epithelium through the back of the throat (retronasal olfaction) as tastants elicit a concurrent taste sensation (**Figure 1A**). Both sensations are then perceived as a holistic flavour identity. Indeed, so strong is the quasi-synaesthetic relationship between taste and odour that retronasal odours not only enhance taste perception^1,2^ but can even elicit a taste percept in the absence of a tastant^3^. For example, For example, strawberry aroma in the absence of sweet receptor stimulation still ‘tastes’ distinctly ‘sweet’. Moreover, words used to describe odours tend to either refer to their source or an associated taste, which implies that odours are conceptually mapped to objects or tastes^4,5^. While verbal representations of odours might overlap with tastes, it remains poorly understood to what extent these canonically parallel pathways overlap in the central nervous system.

**Figure 1.**
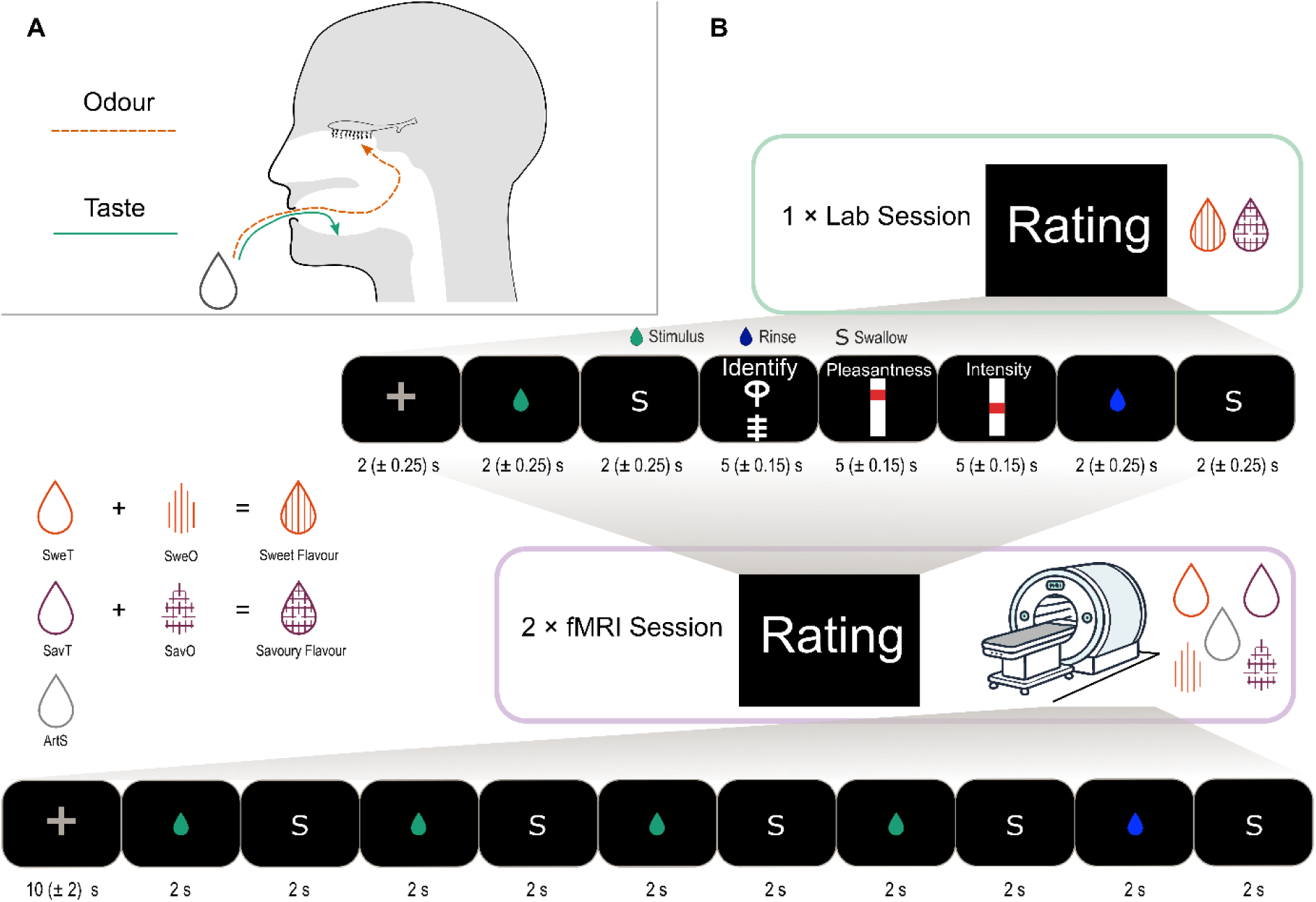
Pathways of chemosensation in the oral cavity and experimental paradigm. **A)** Different routes for taste and odour during food consumption. While the flavour percept is holistic, it relies on chemosensation through parallel transduction routes. **B)** Setup of the experiment. Participants attended 1 laboratory session where they are familiarised with the sweet and savoury flavour mixtures. In two subsequent sessions, they undergo fMRI while receiving unimodal stimuli. SweT: sweet taste; SweO: ‘sweet’ odour; SavT: savoury taste; SavO: ‘savoury’ odour; ArtS: artificial saliva.

The primary sensory cortex involved in the processing of odours has been localised to the piriform cortex in a range of mammalian species. There is a level of preferential processing in the piriform, where the anterior piriform cortex encodes the structure of an odorant and the posterior piriform cortex encodes the identity^6^. Neural encoding of tastants, on the other hand, has highlighted the insula as the primary gustatory cortex, shown in rodents^7^, non-human primates^8^ and humans^9,10^. Curiously, neurons in the gustatory cortex are also known to respond to odours^11–13^. Recent literature has shown reliably distinct ensemble encoding of retronasal odours in the rodent gustatory cortex^14^. Olfactory-induced activations in the insula are unexpected as, canonically, information from primary sensory cortices is processed in parallel before integration by the orbitofrontal cortex (OFC), which encodes the identity^8,15^ and subjective value^15–17^ of a food item. However, gustatory and olfactory stimulation occur almost simultaneously and with great contingency during food consumption, likely encouraging Hebbian pairing between neurons responsive to both chemosensory signals. Both behavioural^3,18,19^ and functional neuroimaging data^20,21^ show the existence of taste-odour associations that imply a strong link between these chemosensory signals. This integration may occur prior to its interaction with other sensory properties of foods in the OFC. In non-human primates, monosynaptic projections from the piriform olfactory cortex to the dysgranular and agranular insula are well-documented^22,23^ and constitute a putative pathway for olfactory-gustatory integration during food flavour perception. We therefore hypothesize that a similar pathway is leveraged in humans, where olfactory information is integrated with gustatory information in the insula to form a holistic flavour concept to induce taste-like activations to odours with prior taste associations. Additionally, ensemble taste representations in the rodent gustatory cortex are known to shift with repeated exposure^24,25^. This key feature, along with encoding of retronasal odours in the insula, may also be preserved in humans.

Here, we use oral delivery of unimodal chemosensory stimulation (tasteless odorants and odourless tastants; **Figure 1A**) and functional magnetic resonance imaging (fMRI) to investigate early integration of chemosensory information in the insula. Specifically, we test if object identity of retronasal taste-associated odours elicit dissociable patterns of activation in the gustatory cortex and if this encoding overlaps with the associated tastes. We show that familiar taste-odour pairs share overlapping encoding patterns in a taste-responsive region of the insula. Crucially, after parcellating the insula based on layer morphology (granularity), we show that this crossmodal overlap is observed mostly in the ventral anterior subregion of the insula, which corresponds to areas of lower granularity. Further analyses to characterise temporal fluctuations in taste and flavour representations showed instability of taste identity patterns, which to our knowledge is the first formal testing of gustatory representational drift in humans.

## RESULTS

### Task design

We tested the pre-registered hypothesis of taste-like activation patterns in response to retronasal odour using fMRI in 25 participants (11 male, 14 female by self-report) while they orally received unimodal taste or odour stimuli. Participants attended a laboratory session where they received congruent flavours (a sweet solution with a ‘sweet’ aroma and a savoury solution with a ‘savoury’ aroma) while performing identification and rating tasks. In two subsequent separate scanning sessions, they underwent fMRI during a passive tasting task where they orally received the unimodal components of the previously exposed flavour stimuli (odourless tastants or tasteless odorants – see **Figure 1B**). Prior to each scanning session, participants also rated the pleasantness and intensity of each stimulus and performed the identification task again to ensure that they were indeed able to differentiate the sweet and savoury stimuli. **Supplementary Figure 1A** shows the distribution of the odours used. As seen in **Supplementary Figures 1B and 1C**, variation in individual-level responses did not affect perceived pleasantness or intensity at the group level.

### Specific sensory cortex activation in response to modality-specific stimulation

We first used mass-univariate General Linear Model (GLM) analyses of modality-specific stimulation, contrasted against the artificial saliva (ArtS) control to test which regions show activation in response to chemosensory stimulation. As seen in **Figure 2A**, oral delivery of tasteless odorants induced whole-brain corrected significant BOLD activations in the left piriform cortex ([-18, -2, -14], z = 5.56, *p_FWE_* = .022), which is part of the primary olfactory cortex. As the piriform cortex was a pre-registered a priori region of interest, we used a small-volume correction (SVC) based on previous statistical localisation by activation likelihood estimation^26^ to show significant activation in the right piriform cortex ([26, 0, -16], z = 4.17, *p_FWE-SVC_* = .005) in response to retronasal odour stimulation.

**Figure 2.**
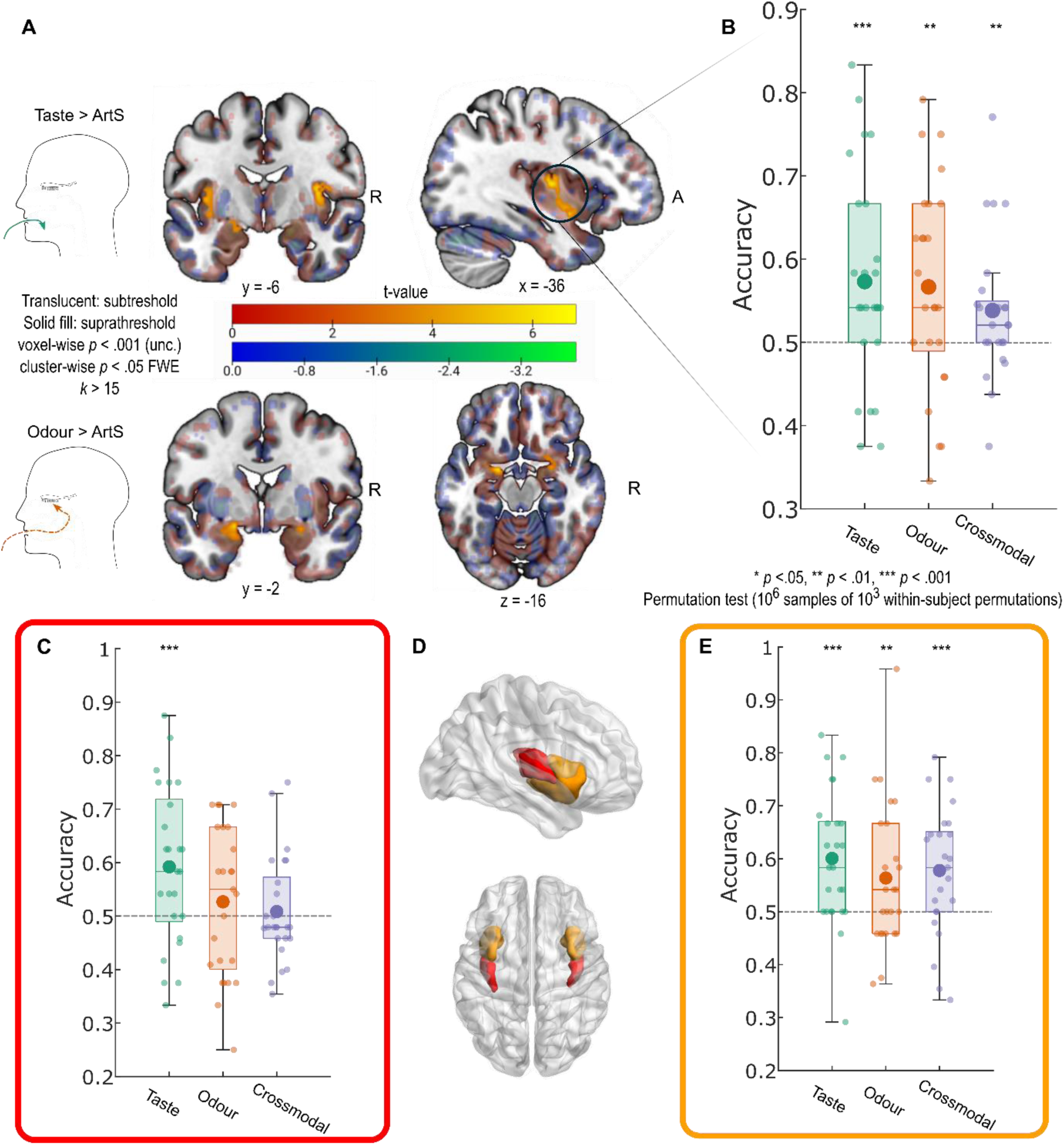
Modality-specific univariate responses and ROI decoding. **A)** *Top*. Univariate contrast of all taste presentations against tasteless artificial saliva (ArtS) shows BOLD response in bilateral dorsal mid-insula extending to ventral anterior insula. *Bottom*. Univariate contrast of all retronasal odour presentations against ArtS shows BOLD response in bilateral piriform cortex. **B)** Leave-one-subject-out taste ROI decoding analyses showing strong dissociability of taste and odour identity, as well as crossmodal decoding between taste and odour. **C)** Significant taste decoding in the dorsal granular insula. **D)** ROIs for dorsal granular insula (red) and dysgranular and agranular insula (orange). **E)** Significant taste, odour and crossmodal decoding in the dysgranular and agranular insula. Large dots signify means; boxes show the interquartile ranges (IQR); whiskers show minimum and maximum ranges using 1.5 IQR. Grey dashed lines signify theoretical chance accuracy. * *p* < .05, ** *p* < .01, *** *p* < .001 permutation test (10^6^ samples of 10^3^ within-subject permutations).

Activations elicited by odourless tastants contrasted against the ArtS control peaked in the bilateral mid-dorsal insula ([-36, -10, 10], z = 5.16, *p_FWE_* = .004; [36, -6, 12], z = 4.66, *p_FWE_* = .018), consistent with activation of canonical primary taste cortex. Notably, the cluster in the left hemisphere extended to parts of the ventral anterior insula (**Figure 2A**), and a significant cluster was seen in the right ventral anterior insula after SVC ([10, 4, -8], z = 4.46, *p_FWE-SVC_* = .010).

### Identity-specific encoding of chemosensory stimulus quality in the insula

In order to test for the presence of identity-specific encoding of chemosensory stimuli in the insula beyond modality-specific univariate activations, we performed multivariate pattern decoding (MVPA) in independent functional ROIs in the bilateral insula responsive to tastants, generated using leave-one-subject-out cross-validation (see **Methods**). We generated a GLM with quality-specific regressors for odours and tastants, i.e., one regressor each for sweet taste (SweT), savoury taste (SavT), sweet odour (SweO), savoury odour (SavO) and artificial saliva (ArtS) and trained a support vector machine (SVM) on the resultant betas from all runs but one (leave-one-run-out cross-validation, see **Methods**). As hypothesised, the classifier could differentiate sweet and savoury taste (mean accuracy of 57.31%, permutation test *p* < .001), confirming that taste-responsive regions in the bilateral insula show dissociable patterns of activation in response to different tastants (**Figure 2B**). Notably, odour quality could similarly be decoded (mean accuracy of 56.67%, permutation test *p* = .003), indicating that this taste-responsive region also shows differential activation to taste-associated odours. We subsequently tested whether encoding patterns of taste-associated odours in the primary gustatory cortex overlap with those of their associated tastes by training a classifier on taste quality and testing it on odour quality and vice versa. Indeed, this crossmodal classifier decoded taste or odour quality significantly above chance (mean accuracy = 53.83%, permutation test *p* = .009), indicating that flavour-specific insular representations of flavour quality are conserved across stimulus modalities. To rule out the possibility that these decoding accuracies were driven by hedonic differences, we examined the voxel-voxel correlation across days against the across-day difference of pleasantness ratings (**Supplementary Figure 2B**) but did not find a significant relationship.

Given that insular projections from the piriform cortex largely terminate in its dysgranular and agranular sections in both primates and rodents^23,27^, we tested for differential identity-specific responsiveness to flavour components in sub-regions parcellating the ROI based on granularity (**Figure 2D**). The granular insula (**Figure 2C**) displayed reliable taste encoding with a mean accuracy of 59.22% (*p* < .001), but no significant encoding of odours (mean accuracy 52.67%, *p* = .127) or crossmodal encoding of flavour identity (mean accuracy = 50.84%, *p* = .308). On the other hand, the dysgranular and agranular anterior insula (**Figure 2E**) displayed population-based encoding of tastant identity (mean accuracy 60.06%, *p* < .001), odour identity (mean accuracy 56.35%, *p* = .004) and crossmodal identity-specific encoding between the two modalities (mean accuracy 57.78%, *p* < .001). These results highlight the insula’s unique role in flavour perception, starting from processing of basic tastant quality in the granular insula and its integration with odour quality to form a flavour percept which is encoded in the dysgranular and agranular insula.

### Whole-brain population-based encoding of chemosensory stimulus quality

In order to investigate whether any region outside the insula displays shared encoding between flavour components, we performed a whole-brain searchlight analysis with our crossmodal MVPA. **Figure 3** shows clusters that survive threshold-free cluster enhancement (TFCE) of 10^4^ permutations. We were able to decode taste from odour and odour from taste in a region of the ventral anterior insula that slightly overlaps but is anterior to the taste-responsive ROI (local peak z = 1.84, *p_TFCE_* = .032), the middle frontal gyrus extending into the lateral orbitofrontal cortex (lOFC; local peak z = 1.88, *p_TFCE_* = .030) and the medial orbitofrontal cortex (mOFC; local peak z = 1.95, *p_TFCE_* = .036). However, the highest crossmodal decoder performance was observed in a region extending from the middle occipital gyrus to the bilateral cuneus (peak z = 2.82, *p_TFCE_* = .002), an area that does not typically form part of the flavour network. As each flavour was assigned an abstract visual stimulus, occipital gyrus decoding accuracy might be attributed to activity related to the conditioned visual cues. Taken together, these results suggest that common representations between flavour components, regardless of modality, exist in various cortical regions beyond established flavour-responsive regions, indicating the presence of a flavour-sensitive cortical network. The fact that the insula is the only canonical primary chemosensory cortex in this network indicates its role as a putative integrating hub.

**Figure 3.**
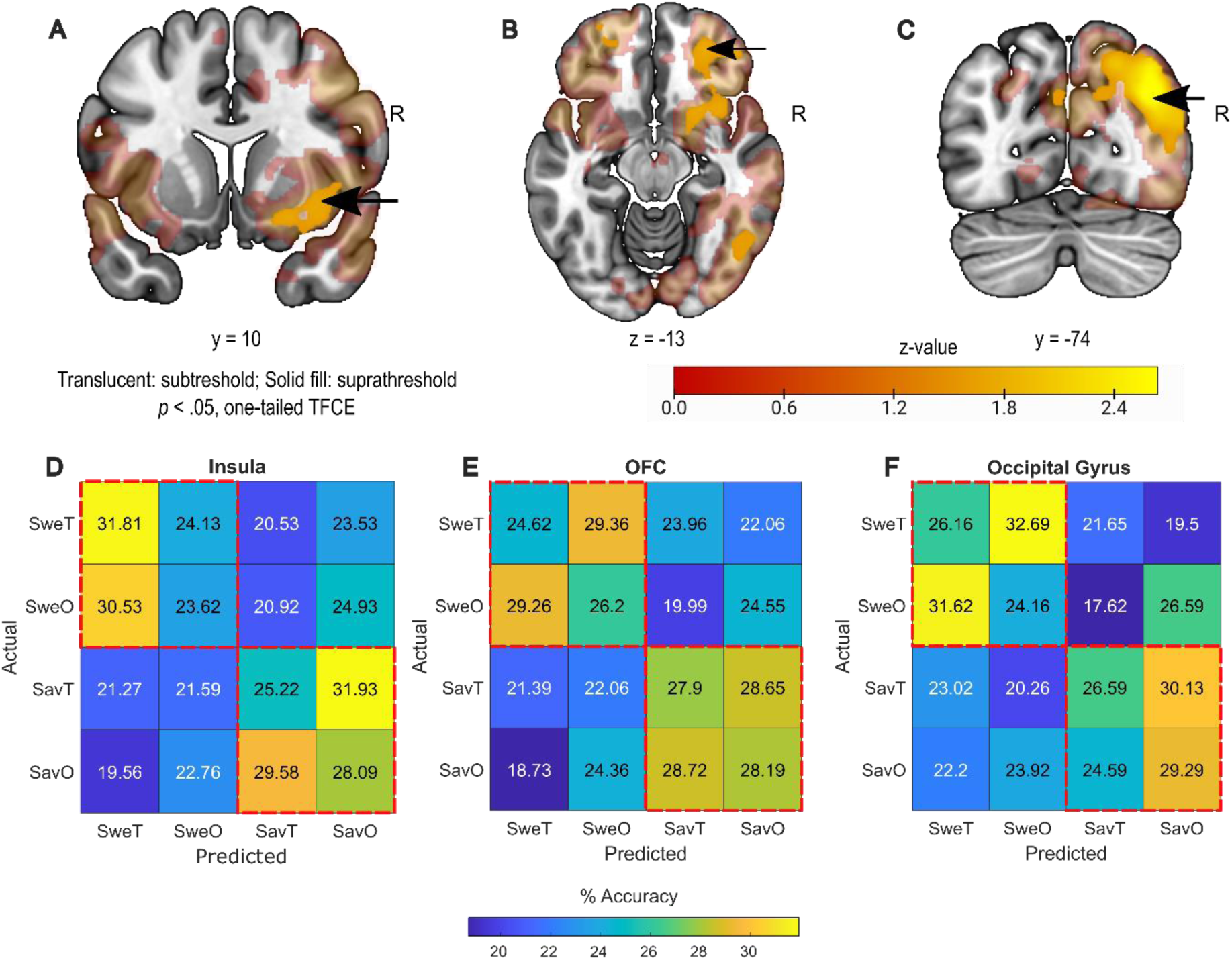
Whole-brain searchlight crossmodal decoding and confusion matrices. **A – C)** Z-map of searchlight decoding accuracy derived from TFCE, showing significant crossmodal decoding accuracy in the ventral anterior insula (A), the lateral orbitofrontal cortex (B) and the middle occipital gyrus (C). **D)** Confusion matrix from the insula peak showing bias for prediction of SweT for both sweet stimuli, whereas prediction for savoury stimuli is not biased towards taste or odour. **E – F)** Confusion matrix from the OFC (E) and occipital gyrus (F) peaks showing no bias towards sweet taste prediction. SweT – Sweet taste; SweO – ‘Sweet’ Odour; SavT – Savoury Taste; SavO – ‘Savoury’ Odour. Red dashed boxes indicate shared flavour identities. Arrows in **A – C** indicate the central voxel of the spherical ROIs used for **D – F**.

For cortical regions with modality-independent chemosensory encoding patterns (ventral anterior insula, lateral OFC and middle occipital gyrus), we proceeded to examine differences in how these patterns were organised. A confusion matrix extracted from a spherical ROI around the insula searchlight peak showed that both sweet tastes and ‘sweet’ odours were more likely to be predicted to be sweet taste, whereas savoury taste and ‘savoury’ odour were confused for each other (**Figure 3D**). Conversely, a confusion matrix extracted from a spherical ROI around the lOFC peak shows that stimuli of the same flavour quality were equally likely to be confused for each other (**Figure 3E**). A similar pattern was also observed for the sweet stimuli in the confusion matrix extracted from a spherical ROI around the occipital gyrus peak, whereas savoury stimuli of both modalities were more likely to be predicted to be the ‘savoury’ odour here (**Figure 3F**). The differences in these confusion matrices imply encoding differences amongst these regions despite shared flavour-specific patterns in response to flavour components. While sweetness and savouriness are part of different flavour identities, savouriness is rarely encountered as a pure taste in the real world, whereas pure sweet tastants are ubiquitous. For example, while most are familiar with the taste of pure sugar, pure savoury tastes (from sources such as aqueous glutamate) are typically found in mixed savoury dishes that involve multiple sensory properties. We therefore propose that participants were more likely to associate the sweet percept (regardless of modality) with a taste-like sensation but savouriness as a holistic flavour concept consisting of both aroma and taste. The bias towards predicting sweet stimuli of either modality as sweet taste in the insula, and the lack thereof with savoury stimuli, highlights the putative role of the insula in integrating and encoding the flavour percept.

### Representational drift of flavours over time

While we have shown overlapping taste and odour identity patterns in the insula, we were interested if these patterns change over time, as shown in the rodent gustatory cortex^24,25^. We therefore proceeded to characterise changes in insular taste encoding patterns over scanning sessions and potential similar temporal declines in odour identity representations. Specifically, we compared data trained and tested from the same scanning session against those trained and tested from different sessions. Confirming previous observations of temporal change in taste identity representations, decoder performance for taste quality dropped significantly when the decoder was trained and tested on different days (**Figure 4A**; Two-way ANOVA F_Train x Test_ (1,92) = 4.21, *p* = .043), indicating that pattern-based encoding of taste qualities in the insula is not stable across days. Of interest, this temporal sensitivity was not observed for the decoder trained and tested on odour quality or crossmodal decoding (**Figure 4B – C**), indicating that insular odour representations are more temporally stable.

**Figure 4.**
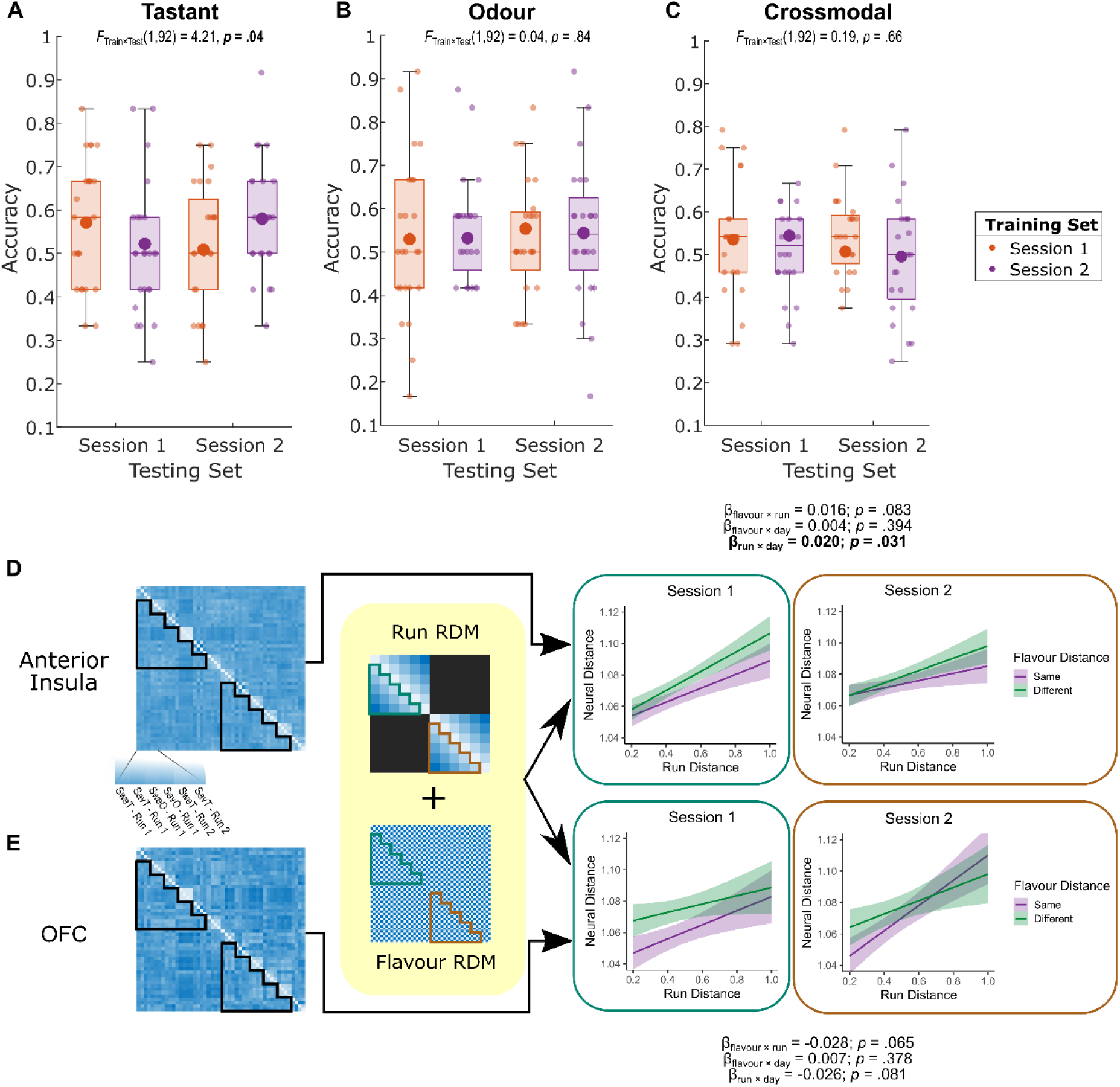
Temporal dynamics of taste, odour and crossmodal neural signals. **A)** Significant differences in accuracy between decoders trained and tested on taste identity within the same day as opposed to across days. **B – C)** No significant difference in within-day and across-day accuracies for decoders trained and tested on odour identity or using crossmodal partitioning. Large dots signify means; boxes show the interquartile ranges (IQR); whiskers show minimum and maximum ranges using 1.5 IQR. **D)** Representational drift of flavour representations in the insula across runs. Model of the neural RDM from betas of all four conditions across runs (left) as predicted by the runwise distance and the flavour distance. Significant reduction in representational drift in the insula between session 1 and session 2. **E)** Same as D but using the neural RDM extracted from the OFC searchlight peak, showing no reduction in drift between days.

While taste encoding specifically appears to show session-related changes, we were also interested in examining if and how flavour representations changed across runs in the same session. To that end, we applied linear mixed-effects modelling (LMM) in Representational Similarity Analysis (RSA) using a novel quantification of the runwise drift in neural representation. We obtained the scaled pairwise distance between each run number, such that the runs furthest apart had a distance of 1 and proximal runs had the smallest distance (see **Figure 4D**). This measure of runwise distance was used as a predictor variable in an LMM in addition to flavour distance to model how well these distances predicted neural dissimilarity between runs (see **Methods**), where distances from the same run were excluded from analysis.

**Figure 4D** shows baseline representational drift (as judged by the representational drift of the same flavour across runs) in both the insula and the OFC. There was greater drift in insular flavour representation in the first session compared to the second session (β_run × day_ = 0.020, *p* = .031) and subthreshold divergence of different flavours (β_flavour × run_ = 0.016, *p* = .083). These differences in drift, both the steeper runwise drift on the first day and the trend towards a divergence, are consistent with stimulus identity encoding in the insula, where representations of different flavours are preserved across runs, particularly in the first session. On the other hand, flavour representational drift in the OFC was not significantly reduced on the second day (**Figure 4E**; β_run × day_ = -0.026, *p* = .081) and had a subthreshold convergence of flavour representation (β_flavour × run_ = -0.028, *p* = .065), indicating that dissimilarity between similar and different flavours was conserved between proximal runs but was less separable between distal runs. This convergence may be indicative of value-level processing in the OFC, as the subjective value of a stimulus is likely to fluctuate over the course of a session, resulting in some preservation of value in proximal runs but a decline in distal runs.

## DISCUSSION

Our study demonstrates that tastes and retronasal odours evoke a shared flavour-specific neural code in the dysgranular and agranular insular cortex, but not in granular insula or piriform cortex. We also use a novel quantification of runwise distance to demonstrate that insular representations of flavours change within the course of a scanning session, although this drift is less prominent in the second session. Taken together, these findings for the first time provide a neural basis for the quasi-synaesthetic perception of the two during food consumption. Our findings highlight the role of the insula as a potential hub for the integration of flavour components. This putative early integration both explains the strong perceptual link between tastes and odours and can be explained by simultaneous stimulation arising from both modalities during food consumption. The fact that odours and taste evoke comparable neural codes build the likely neural foundation for the frequently observed confusion between odour and taste during food consumption, as well as their mutual enhancement.

In this study, both sweet and savoury taste stimulation induced activations in the bilateral dorsal mid-insula extending to the ventral anterior insula, which replicates and reinforces previous literature implicating the insula as the primary gustatory cortex^28,29^. In addition to the expected presence of taste quality coding within this taste-responsive region, which was in line with previous MVPA studies^9,30^, we also note that this region reliably displays pattern-based encoding of taste-associated odour identity. While the anterior insula and frontal operculum have been known to display activation in response to olfactory cues, particularly food-related odours^20,21^, involvement has been less consistently demonstrated for retronasal odour delivery. Here, we demonstrate for the first time that, even if the absence of a robust mass-univariate retronasal odour signal in these regions, that the human gustatory cortex spatially encodes retronasal odour qualities as well as tastes. These findings echo retronasal odour encoding in the rodent gustatory cortex^14,31^, despite the differences in gustatory projections between humans and rodents^32,33^.

Of notable interest, we show that odour identity patterns in the insula overlap with those elicited by their associated tastes. This provides a putative neural basis for odour-induced taste sensation in the absence of a tastant^3^: that is, a taste sensation occurs because the gustatory cortex responds as it would to a real tastant. This phenomenon has not previously been observed in other species, as previous experiments investigating the role of the insula in processing olfactory information have found no response^8,34^, a univariate activation^20,21,35^ or no crossmodal encoding between familiar tastes and odours^14,31^. However, a crucial distinction is that, in this study, congruent taste-odour pairings were used, and participants were familiarised with the taste-odour combination to maximise the flavour binding between the chemosensory flavour components. We propose that this strong flavour binding is key, as a taste-odour association is required for an odour-induced taste sensation, which in turn is reflected by taste-like neural patterns in response to odours.

The insula is a large structure with complex variations in cytoarchitecture, most notably the anteroposterior gradient of granularity observed in primates and humans^36–40^. Given the existence of only five known basic taste qualities^41^, it is unlikely that a structure with such size and complexity would store basic taste representations without further processing, such as integration with other chemosensory properties of foods. Anatomically, the ventral anterior insula has reciprocal projections from olfactory areas such as the piriform cortex and orbitofrontal cortex^22,23,42^, and functional perturbation of these regions induces neuronal firing in the others^43,44^. In our study, spatial encoding of taste identity was spread throughout the granular, dysgranular and agranular portions of the insula, whereas taste-associated odour encoding was restricted to the dysgranular and agranular insula. This distinction, along with the fact that the insular odour encoding patterns overlap with their associated tastes, implies that the flavour percept is integrated in the dysgranular and agranular insula based on gustatory input from the granular insula and olfactory input from the piriform cortex, as previously proposed by Small *et al*. (2013)^45^.

The existence of a common neural code for associated gustatory and olfactory information in the insula implies that chemosensory signals that constitute flavour perception are truly integrated into a shared object representation prior to their value-level processing in the OFC. This mechanism goes against the established role of the OFC as the integrating hub of all sensory modalities related to food reward, given direct afferent OFC projections from visual, auditory, tactile, gustatory and olfactory sensory cortices^15^. Nevertheless, early chemosensory integration in the insula does not necessarily rule out the role of the OFC in evaluating the subjective value of food reward. In humans and other primates, OFC activity does reflect an identity-based value signal in orthonasally presented odours^46–48^ as well as nutrient-guided valuation of visual food stimuli^49,50^. In addition to these anticipatory cues, the human and primate OFC is also sensitive to consummatory reward features such as taste^16,35^, retronasal odour^35,51^ and oral texture^52,53^. Given the established role of OFC neurons in encoding the identity and value of offered and chosen oral food stimuli^17^, as well as our finding of crossmodal decoding in the OFC, our results are in line with the idea that the OFC evaluates an integrated flavour signal from the insula.

Building on the evidence of taste-odour integration in the insula, we next examined how these representations change over time. Our study pioneered formal testing of changes in taste encoding patterns across days in humans. Monocellular and ensemble taste identity representations in the rodent gustatory cortex are known to shift^7,24,25^. Furthermore, while such a change has been indicated in a high-field human fMRI study^9^, no formal testing of within-day and across-day decoding accuracies was conducted. Inspired by these studies, we designed our experiment to allow testing for similar shifts in humans. In doing so, we show that taste encoding does indeed change across days, with the across-day partitioning strategy performing significantly poorer than the within-day partitioning, thereby indicating that ensemble taste patterns in the human insula also change across days, similar to observations in rodents. By using fMRI RSA to characterise run-wise representational drift of flavour identity in the insula and the OFC, we observed that flavour patterns in the insula drift more on the first day compared to the second day, exhibiting slight divergence. On the other hand, there is pattern convergence and a lack of a day-wise change in representational drift in the OFC. Based on these results, we propose that the insular patterns observed encoded identity, whereas the OFC patterns encoded value. In the OFC, representational distance between the same flavour and different flavours was largest in proximal runs but similar in distal runs. This implies that the OFC encodes a feature of the stimuli that differentiates flavours but does not remain stable throughout the session. On the other hand, insular representational distance between the same and different flavours is preserved throughout the session, if exhibiting a slight divergence. This implies that the insula holds representations of a stimulus feature that remains constant throughout the session, such as flavour identity.

The low temporal resolution of fMRI does not allow us to directly test that the flavour signal in the insula occurs prior to one in the OFC. Theoretically, an alternative interpretation might be that taste-like responses in the anterior insula to olfactory stimulation arise due to back-projection after flavour recognition in the OFC^54^. However, our findings combine with primate tractography showing reciprocal monosynaptic projections between the insula and the piriform cortex^23,27,42^ and functional experiments showing early signalling of retronasal odour in the rodent gustatory insular cortex^12–14,31^ to strongly suggest that flavour representation occurs in the insula upstream of the OFC, in addition to providing a foundation for future studies to directly test that hypothesis.

In addition to demonstrating overlapping encoding of odours and their associated tastes in the insula, our study expands on previous evidence of the roles of primary sensory cortices in response to unimodal stimuli. Despite robust univariate activation in the piriform cortex in response to olfactory stimulation, our pattern-based analyses were unable to decode between the two odours in the piriform cortex (**Supplementary Fig 3**). On the other hand, robust decoding of odour quality was possible in the primary gustatory cortex in absence of univariate odour activation. Given prior findings demonstrating spatial encoding of odour identity in the piriform cortex for orthonasal odours^6,55^, and the established role of the piriform cortex in odour processing in a variety of mammals^6,12,26,31,51^, this result was unexpected. One explanation for this divergence may be that the resolution used by our scanning protocol was unable to capture the fine-grained differences in activation patterns between the odour stimuli. While we cannot rule out that decoding would be possible at finer spatial resolution, the fact that odour identity decoding was in fact possible in the insular ROI may also indicate fundamental differences in identity coding for orthonasally and retronasally presented odours. Therefore, future research should consider higher spatial resolutions to resolve this issue.

For the present study, avoiding trial-by-trial ratings, particularly of pleasantness, was a conscious decision based on previous evidence showing task-based variability in insular activation^56^. Tasks involving explicit pleasantness ratings inflate differential activation between taste-containing and tasteless stimuli compared to detection, identification and passive tasting paradigms^57^. Electing for a passive tasting paradigm, similar to Avery *et al*. (2020)^9^, ensured that the observed taste signal was driven by the presence of a tastant, rather than attentional effects. Furthermore, we combined explicit pleasantness ratings collected on both days with voxel-voxel correlation methods in supplementary analyses (**Supplementary Fig 2**), where we do not detect a relationship between session-wise shifts in pleasantness and representational distance in the insula, thus indicating that the difference in pattern-based signals is not driven by changes in pleasantness. Due to the lack of trial-by-trial psychophysical and hedonic ratings, we are, however, unable to authoritatively distinguish between cortical sites where overlapping encoding of food odour and taste is consistent with identity-based versus a value-based encoding. An avenue for future research is to develop a design that would allow the dissociation of these possibilities in a targeted manner.

One natural extension of this work is to investigate potential crossmodal encoding between orthonasal olfaction and gustation. Odour is the only food sensory property that is perceived in the anticipatory and consummatory phase, due to orthonasal and retronasal routes of olfaction^3^. Given the role of odour in encouraging approach or avoidance^58^, crossmodal encoding of gustatory and retronasal olfactory cues in the insula may form part of a mechanism whereby odours acquire taste properties during consumption to modulate appetitive behaviour towards sources of the same odour in the outside world^59^. This mechanism, however, hinges on a common representation of odours and tastes of the same food item, particularly in cortical regions that respond to gustatory information. Rodent gustatory cortex ensemble patterns for retronasal odours do not overlap with tastes^14^, and inactivation of the gustatory cortex inhibits retronasal olfactory learning while maintaining orthonasal olfactory learning^60^. However, rodents are obligate nose breathers^61^, whereas humans can and do breathe through the oral cavity^62^, such that there may be more similarity between retronasal and orthonasal olfaction in humans. Furthermore, although single-neuron recordings in the primate insula show no encoding of orthonasal stimulation^8,34^, odours forming part of a flavour percept through constant contingent presentation with a taste stimulus may induce a gustatory cortical response. Moreover, due to the ensemble nature of taste encoding^41^, single-neuronal recordings may fail to capture taste-related ensemble activity.

Taken together, our study demonstrates the role of the insula as a critical hub for flavour integration through taste-odour convergence. Our crucial finding on the crossmodal overlap between odour encoding patterns and their associated tastes extends previous functional neuroimaging studies in various mammalian species implicating the insula as the primary gustatory cortex^8,10,11,34,63^ and evidence for reciprocal insular-piriform projections established through tractography^23,27,42^ and neuronography^43,44^. This finding both explains observations in behavioural studies of odour-induced taste sensations^3^ and likely forms part of a larger mechanism whereby odours acquire taste properties and associated hedonic values to encourage or discourage more consumption of particular foods. This basic mechanism behind flavour integration has implications for flavour preference acquisition and dietary patterns.

## MATERIALS AND METHODS

### Participants

Twenty-eight participants in total completed the fMRI sessions. Of these, 3 had to be excluded due to excessive signal dropout (1), excessive movement during scanning (1) and nausea (1). Therefore, 25 participants (11 male, 14 female) were used in the analysis, of whom one attended only one scanning session. The mean age of the sample was 27.8 years (SD = 6.0 years), and the mean BMI was 22.8 kg/m^2^ (SD = 2.8 kg/m^2^). To be included, participants had to be between 18 and 45 years old, speak fluent English, have a normal sense of smell (tested using the Sniffin’ Sticks Identification task [cut-off score of 12]^64,65^) and a normal sense of taste (tested using tastant sprays in the mouth), have normal or corrected-to-normal vision, not be pregnant, display no cold/flu symptoms and have no known eating disorder. All participants completed an informed consent sheet before taking part in any screening. All procedures were in accordance with the Declaration of Helsinki and approved by the local ethics committee (Regionala etikprövningsnämnden i Stockholm, Dnr 2021-05138).

### Laboratory session

Participants were trained on the task and the stimuli pairing simultaneously in their pre-scanning behavioural session. At the start of the training session, participants filled out a screening questionnaire and performed the taste and odour screening, during which they identified taste qualities of tastants sprayed in the mouth and the odour identities of common objects using the Sniffin’ Sticks protocol^64,65^. Subsequently, each participant completed a short pretesting task by rating the pleasantness and intensity (both on a visual analogue scale between -5 and 5) of three sweet and three savoury flavour mixtures three times in a randomised order from taste cups. Two flavour mixtures, one sweet and one savoury, matching in mean intensity were chosen for the study. Mixtures with a pleasantness rating below -2 were not considered in order to prevent evoking disgust responses in the participant. These mixtures were subsequently used during a series of identification and rating tasks for the rest of the behavioural session. Therefore, each participant had a bespoke sweet and savoury flavour combination.

Prior to the session, participants were randomly assigned two abstract visual cues (letters from the Phoenician alphabet), one for the savoury flavour and another for the sweet flavour. In each trial of the rating task, they participant was presented 1 ml of a stimulus and was asked to choose which visual cue it corresponded to, such that participants would learn the cue associated with each flavour. This cue association allowed us to examine if participants were able to distinguish the flavours without being primed by words such as ‘sweet’ or ‘savoury’. They subsequently rated it for pleasantness and intensity. Parts of the trial that required participant input (’Identify’, ‘Pleasantness’ and ‘Intensity’) lasted for a maximum of 5 seconds but ended after the participant entered the input. Therefore, each trial ranged between 10 s to 25 s. Participants completed up to 60 trials, although from trial 40 onwards the experiment ended once their cumulative accuracy rate was above 85%, thereby ensuring that the participants could sufficiently distinguish between the two flavour stimuli.

### MRI sessions

Participants were instructed to attend two structural and functional MR sessions, scheduled such that the final MR session was within 10 days of the behavioural session. During the MRI sessions, participants orally received unimodal stimuli – that is, only sweet taste (SweT), savoury taste (SavT), ‘sweet’ odour (SweO) or ‘savoury’ odour (SavO) during the mini-block, as opposed to a flavour combination, in addition to artificial saliva. Each participant had bespoke SweO and SavO stimuli derived from the flavour combinations used in their behavioural laboratory session. Liquid stimuli were orally delivered using custom 3D-printed mouthpieces designed for gustatory stimulation in fMRI contexts^66^ connected to peristaltic pumps.

Each MRI session began with a shortened version of the behavioural task from the laboratory session (24 trials) where they performed the identification and rating tasks in the MR scanner bore, prior to any scanning, using the same visual cues that they were assigned in the laboratory session. Participants were briefed that the flavours they would receive might vary slightly from the ones in the laboratory session, such as in terms of intensity. Only participants who scored over 75% accuracy could continue. All participants achieved this score or higher. Between the rating task and the functional runs, participants also underwent a structural multi-echo MPRAGE scan to facilitate image realignment and normalisation.

Participants completed six functional runs per session. Participants were presented with 0.5 ml of the unimodal stimuli or artificial saliva and asked to swallow. This sequence was repeated four times with the same stimulus to form a mini-block (for a total of 2 ml over 16 seconds) and followed by a rinse block consisting of 1 ml of artificial saliva before moving to the inter-trial interval. Between mini-blocks, participants were presented with a grey fixation cross for 8-12 s. Participants were instructed to swallow only when the swallow cue appeared on the screen. The stimulus order was pseudo-randomised such that every stimulus (including artificial saliva) has 3 repetitions per run and that stimuli were not repeated consecutively more than once, and each run consisted of 15 mini-blocks.

### Stimuli

#### Unimodal stimuli

The sweet taste (SweT) stimulus was 9% w/v sucrose (Sigma Aldrich). The savoury taste (SavT) stimulus was 1% monosodium glutamate (Sigma Aldrich).

‘Sweet’ odorants (SweO) used were:

- Golden syrup (DSM-Firmenich)
- Lychee (DSM-Firmenich)
- Raspberry

‘Savoury’ odorants (SavO) used were:

- Smoky bacon (DSM-Firmenich)
- Chicken (DSM-Firmenich)
- Onion (DSM-Firmenich)

All stimuli were dissolved in artificial saliva (ArtS; 25mM KCl + 2.5mM NaHCO_3_ in distilled water). In addition, ArtS was used as a baseline stimulus and rinse solution in the functional MR task in order to present a baseline condition of oral stimulation without activation of the taste cortex^67^.

Each participant completed a short pretesting task before performing the identification and rating task in the laboratory session. The unimodal odorants used on MRI days were the same as the odorants used in the flavour stimuli in the laboratory sessions. For example, a participant who had sucrose + lychee and MSG + chicken in the laboratory session would have sucrose, MSG, lychee and chicken separately in the MRI sessions.

#### Flavour stimuli

Flavour stimuli were created by combining the unimodal stimuli. In general, tastants (either sucrose or MSG) were added to the unimodal odour stimuli to achieve the same concentration as the unimodal stimuli; that is, the flavour stimuli had the same concentration of the odorant as the odour stimuli and the same concentration of the tastant as the taste stimuli.

#### Visual

Each flavour pairing was associated with a visual stimulus cue to avoid ‘sweet’ and ‘savoury’ labels. The visual stimuli (characters from the Phoenician alphabet) were processed, and 14 were deemed suitable for this purpose. Participants were randomly assigned stimuli (one for the sweet flavour and one for the savoury flavour) before training.

### MRI data acquisition and preprocessing

#### Acquisition

Each run consisted of 228 T2*-weighted BOLD gradient multi-echo echoplanar images (EPI) using a Siemens 3 T Prisma scanner running the syngo MR E11 system equipped with a 64-channel head coil. Fifty-two interleaved axial slices of 1.7 mm thickness were collected with the following parameters: in-plane voxel size = 2 mm × 2 mm; slice thickness = 1.7 mm (distance factor of 15%); echo time (TE) = 42.0 ms; repetition time (TR) = 2000 ms; flip angle (FA) = 90°; field of view (FOV) = 208 mm × 208 mm. For brains that exceeded the field of view, partial brain coverage was used, which included the frontal lobe, temporal lobe and occipital lobe.

Prior to the functional scans, a gradient echo image with two echoes was extracted to generate a field map with the following parameters: 2 mm isotropic voxel; TE1 = 4.92 ms; TE2 = 7.38 ms; FA = 60°; TR = 565 ms; FOV = 208 mm × 208 mm; phase encoding direction: R → L axial.

Furthermore, to enable normalisation to a standardised space, a high-resolution multi-echo T1-weighted MPRAGE structural image was acquired with the following parameters: 1 mm isotropic resolution sagittal slices; TE1 = 1.69 ms; TE2 = 3.55 ms; TE3 = 5.41 ms; TE4 = 7.27 ms; FA = 7.0°; acquisition time = 2530 s; FOV = 256 mm × 256 mm. Extracting the root-mean-square (RMS) of the multiple echoes resulted in a high-quality image that was used in subsequent steps.

#### Preprocessing

Images were preprocessed using SPM12 (Wellcome Department of Imaging Neuroscience, Institute of Neurology, London, UK) implemented in MATLAB 2021b (Mathworks). Firstly, a field map for each MRI session was calculated using the gradient echo images, specifically the phase images and the magnitude image, using the in-built SPM12 Field Map Toolbox. Random EPI images from the session were then loaded and unwarped to visually check the extent of distortion correction. The structural images from both MRI sessions were coregistered and a mean image was extracted in order to further improve spatial resolution. This mean structural image was then segmented into tissue probability maps (TPMs), and a skull-stripped brain consisting of the grey matter, white matter and cerebrospinal fluid was calculated (with a combined TPM cutoff of 0.8).

After slice-time interpolation to the 0.99-second slice, functional images from each run were realigned to the first image and unwarped using the previously calculated field map for the pertinent session. They were then coregistered to the skullstripped brain and normalised to MNI space using the deformation field obtained from the segmentation phase. The resultant normalised images were then smoothed with a 6 mm full-width-half-maximum (FWHM) isotropic Gaussian kernel. Finally, both the smoothed and unsmoothed normalised images from each run were detrended using Linear Model of Global Signal detrending^68^ to remove global effects from the time series on top of the pre-registered preprocessing pipeline. These detrended images were used in further analyses.

### Statistical analyses

#### Mass-univariate GLM analyses

First-level general linear models (GLMs) were conducted within each participant using the smoothed and normalised individual BOLD data in SPM12. Within each individual subject, we modelled the following regressors of interest (using a boxcar function with a duration of 16 s):

- All tastant presentations (both sweet and savoury – Condition 1)
- All odorant presentations (both ‘sweet’ and ‘savoury’ – Condition 2)
- All ArtS presentations as the explicit baseline condition (Condition 3)

We also modelled the rinse block for a period of 4 s as a regressor of no interest. The boxcar regressors were convolved with the canonical haemodynamic response basis function (HRF). Furthermore, confound regressors included the six rigid-body motion parameters estimated from the realignment procedure, in addition to spike regressors to censor frames with framewise displacement greater than 1 mm (as calculated from the motion parameters). Runs with more than 10% of their frames censored were excluded from the analysis. A grey-matter explicit mask was used (SPM grey matter TPM thresholded at 0.2) to limit the analysis to only voxels containing grey matter. We also specified the model to include a high-pass filter (HPF) of 128 s to remove slow signal drift.

Within each subject, a contrast of interest (Condition 1 – Condition 3 in the case of tastants and Condition 2 – Condition 3 in the case of odorants) was calculated. The group-level analysis using a one-sample t-test on the globally pooled contrast of interest using an unweighted summary statistics approach. Analyses were conducted using a whole-brain grey-matter mask. Cluster-wise significance was conducted using a threshold of *P_FWE_* < .05, a cluster-cutting threshold of *k* > 15 and voxel-wise threshold of uncorrected *P* < .001. Small-volume corrections (SVCs) were performed from pre-registered peaks, limiting them to a 10 mm radius spherical Region of Interest (ROI). Piriform SVC coordinates were taken from a statistical localisation of the olfactory cortex^26^, whereas anterior insula SVC coordinates were taken from a high-field study on taste activation^9^. ROIs for small-volume corrections and the analysis steps were pre-registered, although we applied a stricter cluster-cutting threshold than pre-registered.

#### Multivariate pattern analysis

Multivariate Pattern Analysis (MVPA) was conducted using the CoSMoMVPA toolbox implemented in MATLAB 2021b. For ROI analyses, the ROIs were formed from functional clusters of the contrast of tastants against ArtS in the mass-univariate GLM analysis to isolate regions that were responsive to taste. In order to avoid bias by using the same participant for ROI generation and analysis, ROIs for each subject were generated from the data of the remaining subjects (leave-one-subject-out cross-validation), using a more lenient voxel-wise threshold of uncorrected *P* < .01 and a cluster-cutting threshold of *k* > 150. Post-hoc ROI MVPA used subregions of the insula as parcellated by Fan et al. (2016).

Prior to any MVPA, first-level GLM analyses were conducted on unsmoothed normalised functional data in order to preserve voxel-level differences in activation. Within each subject, we used the following regressors of interest (using a boxcar function with a duration of 16 s):

- Sweet taste presentation (SweT)
- Savoury taste presentation (SavT)
- ‘Sweet’ odour presentation (SweO)
- ‘Savoury’ odour presentation (SavO)
- Artificial saliva explicit baseline presentation (ArtS)

In addition, rinse periods were modelled as regressors of no interest for a period of 4 s. Regressors were convolved with the canonical HRF basis and confound regressors were the same as the univariate analyses. Similar to the univariate analyses, we also included an HPF of 128 s and applied a whole-brain grey-matter mask.

MVPA was conducted on the resultant runwise betas of the above model. As pre-registered, we trained a linear support vector machine (SVM; implemented in the LIBSVM package^70^) decoder on the beta weights for each voxel obtained from the first-level GLM. We employed a leave-one-run-out cross-validation partition: in each decoding step, the decoder is trained on all runs bar one, and its performance is subsequently tested on the left-out run. For crossmodal decoding analyses, we subtracted the training data mean from both the training and testing data to remove mass univariate differences in activation between different modalities. Whole-brain crossmodal searchlight MVPA used the unsmoothed detrended functional scans in MNI space. At each spherical searchlight with a radius of 4 voxels (8 mm), we applied the same partitioning and mean-centring strategy as the ROI analysis to train and test the decoder. Only grey-matter voxels were used to create the searchlights. We then mapped the average accuracy of the decoder onto the centre voxel of the searchlight before moving onto the next searchlight. We subtracted the theoretical chance level from the accuracy maps before then smoothing them using a 6mm FWHM isotropic Gaussian kernel for group-level analysis.

#### Permutation-based significance testing

Significance testing of classification accuracies was performed via nonparametric permutation testing instead of the pre-registered parametric t-tests against the hypothetical chance level. We elected to perform permutation testing to establish a data-driven null distribution that would account for potential autocorrelations or biases that might lead to differences between a theoretical and a data-driven null distribution ^71,72^. Specifically, in ROI MVPA analyses, the null distributions were created through 10^6^ samples of 10^3^ within-subject permutations. That is, we shuffled the labels (SweT, SavT, SweO, SavO) within each subject and trained and tested the decoder following the same partitioning parameters 10^3^ times. From these within-subject null distributions, we then sampled one accuracy per subject and obtained the group mean accuracy 10^6^ times with replacement. We then calculated one-tailed *P*-values based on the ratio of permuted group mean accuracies higher than the real group mean accuracy.

For whole-brain searchlight analyses, the theoretical chance level (one divided by the number of labels used in the testing set) was subtracted from the accuracy maps from searchlight analyses. These maps were then smoothed and masked such that only grey-matter voxels were used. Finally, we performed prevalence testing on the group data using 10^4^ Monte-Carlo simulations and threshold-free cluster enhancement to account for multiple comparisons^73^ implemented in CoSMoMVPA. Clusters of more than 15 contiguous voxels with a z-score of greater than 1.65 (for one-tailed testing) survived.

#### Representational drift analysis

In order to analyse flavour representational drift across runs, we employed a combination of representational similarity analysis with linear mixed-effects modelling and a novel measure of runwise distance. Specifically, we quantified the runwise distance as the pairwise distance between runs in the same session while excluding data from separate sessions (**Figure 4D – E**), scaled by dividing the distances with the largest runwise distance. Parallel to this, we extracted the betas for each of the four conditions in each run before creating a representational dissimilarity matrix (RDM) by extracting the pairwise correlational distance between them (defined as one minus the voxel-voxel Pearson correlation coefficient). We then designed a linear mixed-effects model with where the neural RDM is predicted by the runwise RDM, the flavour RDM (0 for the same flavour; 1 for different flavours) and the session indicator, as well as their interactions, with the subject identity as the random effect and random intercepts and random slopes for the runwise RDM and the flavour RDM. Linear mixed-effects models were conducted using the inbuilt fitlme function on MATLAB.

#### Data availability and replicability

The design, preprocessing and univariate and taste-responsive ROI MVPA decoding (including across-day MVPA) analyses were pre-registered, along with ROIs (https://osf.io/a8mte). MVPA on the parcellated insular ROI and representational drift analysis across runs were exploratory. Raw individual-level data cannot be shared publicly but will be made available on reasonable request. MATLAB code is available on OSF (DOI 10.17605/OSF.IO/2KRYV).

## Acknowledgements

The authors would like to thank the staff at the Stockholm University Brain Imaging Centre for their help and support in preparing the scanning protocols; DSM-Firmenich and Prof. Thomas Hummel for providing the tasteless aromas used in the study; Dr Christoph Pfeiffer for help with the 3D-printing of the mouthpiece; Zahra Hejazi, Sümeyra Nur Doğan, Hanne Helming and Hilda Lindén for scanning and lab assistance; Dr Gustav Nilsonne for help with pre-registration and Open Science efforts; Leonie Seidel, Dr Anna Gerlicher and members of the Perception Lab for helpful comments and discussions. For Open Access, the authors apply a CC-BY-NC-ND 4.0 licence on any Author Accepted Manuscript arising from this submission.

## REFERENCES

1. Amsellem, S., and Ohla, K. (2016). Perceived Odor–Taste Congruence Influences Intensity and Pleasantness Differently. Chem. Senses 41, 677–684. 10.1093/CHEMSE/BJW078.

2. Small, D.M., and Prescott, J. (2005). Odor/taste integration and the perception of flavor. Exp. Brain Res. 166, 345–357. 10.1007/s00221-005-2376-9.

3. Rozin, P. (1982). “Taste-smell confusions” and the duality of the olfactory sense. Percept. Psychophys. 31, 397–401. 10.3758/BF03202667.

4. Lynott, D., Connell, L., Brysbaert, M., Brand, J., and Carney, J. (2020). The Lancaster Sensorimotor Norms: multidimensional measures of perceptual and action strength for 40,000 English words. Behav. Res. Methods 52, 1271–1291. 10.3758/s13428-019-01316-z.

5. Majid, A., and Burenhult, N. (2014). Odors are expressible in language, as long as you speak the right language. Cognition 130, 266–270. 10.1016/j.cognition.2013.11.004.

6. Howard, J.D., Plailly, J., Grueschow, M., Haynes, J.-D., and Gottfried, J.A. (2009). Odor quality coding and categorization in human posterior piriform cortex. Nat. Neurosci. 12, 932–938. 10.1038/nn.2324.

7. Katz, D.B., Simon, S.A., and Nicolelis, M.A.L. (2001). Dynamic and Multimodal Responses of Gustatory Cortical Neurons in Awake Rats. J. Neurosci. 21, 4478–4489. 10.1523/JNEUROSCI.21-12-04478.2001.

8. Kadohisa, M., Rolls, E.T., and Verhagen, J.V. (2005). Neuronal Representations of Stimuli in the Mouth: The Primate Insular Taste Cortex, Orbitofrontal Cortex and Amygdala. Chem. Senses 30, 401–419. 10.1093/chemse/bji036.

9. Avery, J.A., Liu, A.G., Ingeholm, J.E., Riddell, C.D., Gotts, S.J., and Martin, A. (2020). Taste quality representation in the human brain. J. Neurosci. 40, 1042–1052. 10.1523/JNEUROSCI.1751-19.2019.

10. de Araujo, I.E.T., and Simon, S.A. (2009). The gustatory cortex and multisensory integration. Int. J. Obes. 33, S34–S43. 10.1038/ijo.2009.70.

11. de Araujo, I.E.T., Rolls, E.T., Kringelbach, M.L., McGlone, F., and Phillips, N. (2003). Taste-olfactory convergence, and the representation of the pleasantness of flavour, in the human brain. Eur. J. Neurosci. 18, 2059–2068. 10.1046/j.1460-9568.2003.02915.x.

12. Maier, J.X. (2017). Single-neuron responses to intraoral delivery of odor solutions in primary olfactory and gustatory cortex. J. Neurophysiol. 117, 1293–1304. 10.1152/jn.00802.2016.

13. Maier, J.X., Blankenship, M.L., Li, J.X., and Katz, D.B. (2015). A Multisensory Network for Olfactory Processing. Curr. Biol. 25, 2642–2650. 10.1016/j.cub.2015.08.060.

14. Stocke, S., and Samuelsen, C.L. (2024). Multisensory Integration Underlies the Distinct Representation of Odor–Taste Mixtures in the Gustatory Cortex of Behaving Rats. J. Neurosci. 44. 10.1523/JNEUROSCI.0071-24.2024.

15. Rolls, E.T., and Grabenhorst, F. (2008). The orbitofrontal cortex and beyond: From affect to decision-making. Prog. Neurobiol. 86, 216–244. 10.1016/j.pneurobio.2008.09.001.

16. Grabenhorst, F., Rolls, E.T., and Bilderbeck, A. (2008). How cognition modulates affective responses to taste and flavor: Top-down influences on the orbitofrontal and pregenual cingulate cortices. Cereb. Cortex 18, 1549–1559. 10.1093/cercor/bhm185.

17. Padoa-Schioppa, C., and Assad, J.A. (2006). Neurons in the orbitofrontal cortex encode economic value. Nature 441, 223–226. 10.1038/nature04676.

18. Fondberg, R., Lundström, J.N., Blöchl, M., Olsson, M.J., and Seubert, J. (2018). Multisensory flavor perception: The relationship between congruency, pleasantness, and odor referral to the mouth. Appetite 125, 244–252. 10.1016/j.appet.2018.02.012.

19. Khorisantono, P.A., Fondberg, R., Lundström, J.N., and Seubert, J. (2024). Dissociable effects of hunger, exposure and sensory overlap on flavour liking. 10.31234/OSF.IO/UCBN4.

20. Seubert, J., Ohla, K., Yokomukai, Y., Kellermann, T., and Lundström, J.N. (2015). Superadditive opercular activation to food flavor is mediated by enhanced temporal and limbic coupling. Hum. Brain Mapp. 36, 1662–1676. 10.1002/hbm.22728.

21. Veldhuizen, M.G., Nachtigal, D., Teulings, L., Gitelman, D.R., and Small, D.M. (2010). The Insular Taste Cortex Contributes to Odor Quality Coding. Front. Hum. Neurosci. 4. 10.3389/fnhum.2010.00058.

22. Carmichael, S.T., Clugnet, M.-C., and Price, J.L. (1994). Central olfactory connections in the macaque monkey. J. Comp. Neurol. 346, 403–434. 10.1002/cne.903460306.

23. Mufson, E.J., and Mesulam, M.-M. -Marsel (1982). Insula of the old world monkey. II: Afferent cortical input and comments on the claustrum. 212, 23–37. 10.1002/cne.902120103.

24. Flores, V.L., Tanner, B., Katz, D.B., and Lin, J.-Y. (2022). Cortical taste processing evolves through benign taste exposures. Behav. Neurosci. 136, 182–194. 10.1037/bne0000504.

25. Fontanini, A., and Katz, D.B. (2006). State-Dependent Modulation of Time-Varying Gustatory Responses. J. Neurophysiol. 96, 3183–3193. 10.1152/jn.00804.2006.

26. Seubert, J., Freiherr, J., Djordjevic, J., and Lundström, J.N. (2013). Statistical localization of human olfactory cortex. NeuroImage 66, 333–342. 10.1016/j.neuroimage.2012.10.030.

27. Pritchard, T.C., and Di Lorenzo, P.M. (2015). Central Taste Anatomy and Physiology of Rodents and Primates. In Handbook of Olfaction and Gustation (John Wiley & Sons, Inc), pp. 701–726. 10.1002/9781118971758.ch32.

28. de Araujo, I.E.T., Geha, P., and Small, D.M. (2012). Orosensory and Homeostatic Functions of the Insular Taste Cortex. Chemosens. Percept. 5, 64–79. 10.1007/s12078-012-9117-9.

29. Mazzola, L., Royet, J.-P., Catenoix, H., Montavont, A., Isnard, J., and Mauguière, F. (2017). Gustatory and olfactory responses to stimulation of the human insula. Ann. Neurol. 82, 360–370. 10.1002/ana.25010.

30. Chikazoe, J., Lee, D.H., Kriegeskorte, N., and Anderson, A.K. (2019). Distinct representations of basic taste qualities in human gustatory cortex. Nat. Commun. 10, 1–8. 10.1038/s41467-019-08857-z.

31. Samuelsen, C.L., and Fontanini, A. (2017). Processing of Intraoral Olfactory and Gustatory Signals in the Gustatory Cortex of Awake Rats. J. Neurosci. Off. J. Soc. Neurosci. 37, 244–257. 10.1523/JNEUROSCI.1926-16.2016.

32. Rolls, E.T. (2016). Functions of the anterior insula in taste, autonomic, and related functions. Brain Cogn. 110, 4–19. 10.1016/j.bandc.2015.07.002.

33. Small, D.M., and Scott, T.R. (2009). What happens to the pontine processing?: Repercussions of interspecies differences in pontine taste representation for tasting and feeding. Ann. N. Y. Acad. Sci. 1170, 343–346. 10.1111/j.1749-6632.2009.03918.x.

34. Verhagen, J.V., Kadohisa, M., and Rolls, E.T. (2004). Primate insular/opercular taste cortex: Neuronal representations of the viscosity, fat texture, grittiness, temperature, and taste of foods. J. Neurophysiol. 92, 1685–1699. 10.1152/jn.00321.2004.

35. Small, D.M., Voss, J., Mak, Y.E., Simmons, K.B., Parrish, T., and Gitelman, D. (2004). Experience-Dependent Neural Integration of Taste and Smell in the Human Brain. J. Neurophysiol. 92, 1892–1903. 10.1152/jn.00050.2004.

36. Gallay, D.S., Gallay, M.N., Jeanmonod, D., Rouiller, E.M., and Morel, A. (2012). The Insula of Reil Revisited: Multiarchitectonic Organization in Macaque Monkeys. Cereb. Cortex N. Y. NY 22, 175–190. 10.1093/cercor/bhr104.

37. Mesulam, M.-M., and Mufson, E.J. (1982). Insula of the old world monkey. Architectonics in the insulo-orbito-temporal component of the paralimbic brain. J. Comp. Neurol. 212, 1–22. 10.1002/cne.902120102.

38. Morel, A., Gallay, M.N., Baechler, A., Wyss, M., and Gallay, D.S. (2013). The human insula: Architectonic organization and postmortem MRI registration. Neuroscience 236, 117–135. 10.1016/j.neuroscience.2012.12.076.

39. Pandya, D.N., Van Hoesen, G.W., and Mesulam, M.M. (1981). Efferent connections of the cingulate gyrus in the rhesus monkey. Exp. Brain Res. 42, 319–330. 10.1007/BF00237497.

40. Roberts, T.S., and Akert, K. (1963). Insular and opercular cortex and its thalamic projection in Macaca mulatta. Schweiz. Arch. Neurol. Neurochir. Psychiatr. Arch. Suisses Neurol. Neurochir. Psychiatr. 92, 1–43.

41. Ohla, K., Yoshida, R., Roper, S.D., Di Lorenzo, P.M., Victor, J.D., Boughter, J.D., Fletcher, M., Katz, D.B., and Chaudhari, N. (2019). Recognizing Taste: Coding Patterns Along the Neural Axis in Mammals. Chem. Senses 44, 237–247. 10.1093/chemse/bjz013.

42. Mesulam, M.-M., and Mufson, E.J. (1982). Insula of the old world monkey. III: Efferent cortical output and comments on function. J. Comp. Neurol. 212, 38–52. 10.1002/cne.902120104.

43. Pribram, K.H., Lennox, M.A., and Dunsmore, R.H. (1950). Some connections of the orbito-fronto-temporal, limbic and hippocampal areas of macaca mulatta. J. Neurophysiol. 13, 127–135. 10.1152/jn.1950.13.2.127.

44. Pribram, K.H., and MacLean, P.D. (1953). Neuronographic analysis of medial and basal cerebral cortex. ii. monkey. J. Neurophysiol. 16, 324–340. 10.1152/jn.1953.16.3.324.

45. Small, D.M., Veldhuizen, M.G., and Green, B. (2013). Sensory Neuroscience: Taste Responses in Primary Olfactory Cortex. Curr. Biol. 23, R157–R159. 10.1016/j.cub.2012.12.036.

46. Critchley, H.D., and Rolls, E.T. (1996). Olfactory neuronal responses in the primate orbitofrontal cortex: Analysis in an olfactory discrimination task. J. Neurophysiol. 75, 1659–1672. 10.1152/jn.1996.75.4.1659.

47. Gottfried, J.A., O’Doherty, J., and Dolan, R.J. (2003). Encoding predictive reward value in human amygdala and orbitofrontal cortex. Science 301, 1104–1107. 10.1126/science.1087919.

48. Howard, J.D., Gottfried, J.A., Tobler, P.N., and Kahnt, T. (2015). Identity-specific coding of future rewards in the human orbitofrontal cortex. Proc. Natl. Acad. Sci. U. S. A. 112, 5195–5200. 10.1073/pnas.1503550112.

49. DiFeliceantonio, A.G., Coppin, G., Rigoux, L., Edwin Thanarajah, S., Dagher, A., Tittgemeyer, M., and Small, D.M. (2018). Supra-Additive Effects of Combining Fat and Carbohydrate on Food Reward. Cell Metab. 28, 33–44.e3. 10.1016/J.CMET.2018.05.018.

50. Suzuki, S., Cross, L., and O’Doherty, J.P. (2017). Elucidating the underlying components of food valuation in the human orbitofrontal cortex. Nat. Neurosci. 20, 1780–1786. 10.1038/s41593-017-0008-x.

51. Small, D.M., Veldhuizen, M.G., Felsted, J., Mak, Y.E., and McGlone, F. (2008). Separable Substrates for Anticipatory and Consummatory Food Chemosensation. Neuron 57, 786–797. 10.1016/j.neuron.2008.01.021.

52. Khorisantono, P.A., Huang 黃飛揚, F.Y., Sutcliffe, M.P.F., Fletcher, P.C., Farooqi, I.S., and Grabenhorst, F. (2023). A Neural Mechanism in the Human Orbitofrontal Cortex for Preferring High-Fat Foods Based on Oral Texture. J. Neurosci. 43, 8000–8017. 10.1523/JNEUROSCI.1473-23.2023.

53. Rolls, E.T., Mills, T., Norton, A.B., Lazidis, A., and Norton, I.T. (2018). The Neuronal Encoding of Oral Fat by the Coefficient of Sliding Friction in the Cerebral Cortex and Amygdala. Cereb. Cortex 28, 4080–4089. 10.1093/cercor/bhy213.

54. Rolls, E.T. (2019). Chapter 7 - Taste and smell processing in the brain. In Handbook of Clinical Neurology Smell and Taste., R. L. Doty, ed. (Elsevier), pp. 97–118. 10.1016/B978-0-444-63855-7.00007-1.

55. Raithel, C.U., Miller, A.J., Epstein, R.A., Kahnt, T., and Gottfried, J.A. (2023). Recruitment of grid-like responses in human entorhinal and piriform cortices by odor landmark-based navigation. Curr. Biol. 33, 3561–3570.e4. 10.1016/j.cub.2023.06.087.

56. Veldhuizen, M.G., Bender, G., Constable, R.T., and Small, D.M. (2007). Trying to Detect Taste in a Tasteless Solution: Modulation of Early Gustatory Cortex by Attention to Taste. Chem. Senses 32, 569–581. 10.1093/chemse/bjm025.

57. Bender, G., Veldhuizen, M.G., Meltzer, J.A., Gitelman, D.R., and Small, D.M. (2009). Neural correlates of evaluative compared to passive tasting. Eur. J. Neurosci. 30, 327–338. 10.1111/j.1460-9568.2009.06819.x.

58. Iravani, B., Schaefer, M., Wilson, D.A., Arshamian, A., and Lundström, J.N. (2021). The human olfactory bulb processes odor valence representation and cues motor avoidance behavior. Proc. Natl. Acad. Sci. 118, e2101209118. 10.1073/pnas.2101209118.

59. Khorisantono, P.A., and Seubert, J. (2024). How Are Food Preferences Formed and Changed? Sensory Contributions to Anticipatory and Consummatory Processing of Food Reward. In Smell, Taste, Eat: The Role of the Chemical Senses in Eating Behaviour, L. D. Stafford, ed. (Springer International Publishing), pp. 75–90. 10.1007/978-3-031-41375-9_5.

60. Blankenship, M.L., Grigorova, M., Katz, D.B., and Maier, J.X. (2019). Retronasal Odor Perception Requires Taste Cortex, but Orthonasal Does Not. Curr. Biol. 29, 62–69.e3. 10.1016/j.cub.2018.11.011.

61. Harkema, J.R., and Morgan, K.T. (1996). Normal Morphology of the Nasal Passages in Laboratory Rodents. In Respiratory System, T. C. Jones, D. L. Dungworth, and U. Mohr, eds. (Springer), pp. 3–17. 10.1007/978-3-642-61042-4_1.

62. Ward, K.A., Nicholls, D.P., and Stanford, C.F. (1993). The prevalence of preferential nasal breathing in adults. Respir. Med. 87, 295–297. 10.1016/0954-6111(93)90026-v.

63. Lemon, C.H., and Katz, D.B. (2007). The neural processing of taste. BMC Neurosci. 8, 1–8. 10.1186/1471-2202-8-S3-S5/FIGURES/2.

64. Hummel, T., Sekinger, B., Wolf, S.R., Pauli, E., and Kobal, G. (1997). ‘Sniffin’ Sticks’: Olfactory Performance Assessed by the Combined Testing of Odor Identification, Odor Discrimination and Olfactory Threshold. Chem. Senses 22, 39–52. 10.1093/CHEMSE/22.1.39.

65. Oleszkiewicz, A., Schriever, V.A., Croy, I., Hähner, A., and Hummel, T. (2019). Updated Sniffin’ Sticks normative data based on an extended sample of 9139 subjects. Eur. Arch. Otorhinolaryngol. 276, 719–728. 10.1007/S00405-018-5248-1/TABLES/2.

66. Munoz Tord, D., Coppin, G., Pool, E.R., Mermoud, C., Pataky, Z., Sander, D., and Delplanque, S. (2021). 3D-Printed Pacifier-Shaped Mouthpiece for fMRI-Compatible Gustometers. eNeuro 8. 10.1523/ENEURO.0208-21.2021.

67. de Araujo, I.E.T., Kringelbach, M.L., Rolls, E.T., and McGlone, F. (2003). Human cortical responses to water in the mouth, and the effects of thirst. J. Neurophysiol. 90, 1865–1876. 10.1152/jn.00297.2003.

68. Macey, P.M., Macey, K.E., Kumar, R., and Harper, R.M. (2004). A method for removal of global effects from fMRI time series. NeuroImage 22, 360–366. 10.1016/j.neuroimage.2003.12.042.

69. Fan, L., Li, H., Zhuo, J., Zhang, Y., Wang, J., Chen, L., Yang, Z., Chu, C., Xie, S., Laird, A.R., et al. (2016). The Human Brainnetome Atlas: A New Brain Atlas Based on Connectional Architecture. Cereb. Cortex N. Y. NY 26, 3508–3526. 10.1093/cercor/bhw157.

70. LIBSVM: A library for support vector machines: ACM Transactions on Intelligent Systems and Technology: Vol 2, No 3 https://dl.acm.org/doi/10.1145/1961189.1961199.

71. Stelzer, J., Chen, Y., and Turner, R. (2013). Statistical inference and multiple testing correction in classification-based multi-voxel pattern analysis (MVPA): random permutations and cluster size control. NeuroImage 65, 69–82. 10.1016/j.neuroimage.2012.09.063.

72. Valente, G., Castellanos, A.L., Hausfeld, L., De Martino, F., and Formisano, E. (2021). Cross-validation and permutations in MVPA: Validity of permutation strategies and power of cross-validation schemes. NeuroImage 238, 118145. 10.1016/j.neuroimage.2021.118145.

73. Smith, S.M., and Nichols, T.E. (2009). Threshold-free cluster enhancement: addressing problems of smoothing, threshold dependence and localisation in cluster inference. NeuroImage 44, 83–98. 10.1016/j.neuroimage.2008.03.061.

