## Supplementary Information for "Tastes and retronasal odours evoke a shared flavour-specific neural code in the human insula"

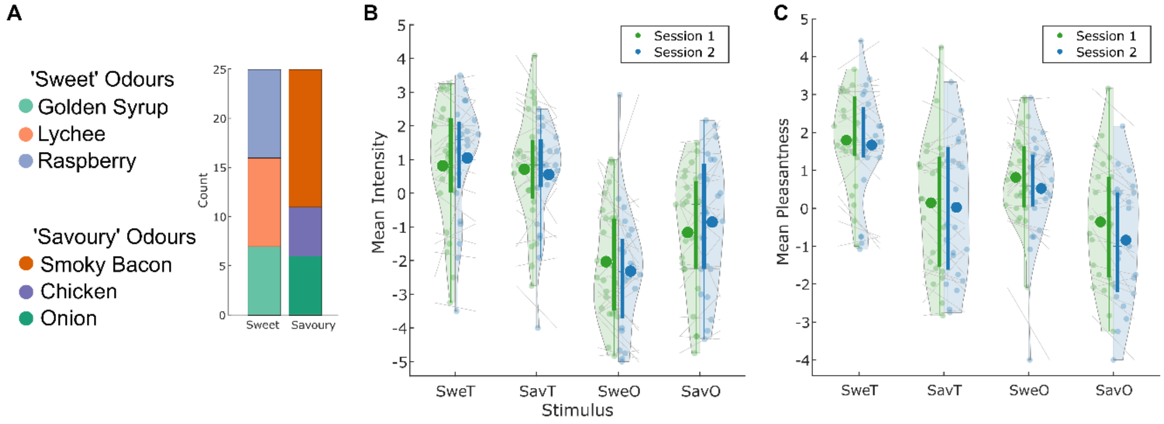


**Supplementary Figure 1.** Distribution of odours and behavioural ratings. **A)** Distribution of the number of participants assigned to the various 'sweet' and 'savoury' odours showing roughly equal split among 'sweet' odours. For the majority of participants, Smoky Bacon was used as the 'savoury' odour. **B)** Group-level and participant-level intensity ratings by session. **C)** Group-level and participant-level pleasantness ratings by session. While individual ratings changed, mean ratings remained largely consistent at the group-level for both pleasantness and intensity. Large dots signify means; small dots signify individual means; box limits indicate IQR; whiskers signify range as 1.5 times IQR; grey lines indicate subject-level changes across sessions.

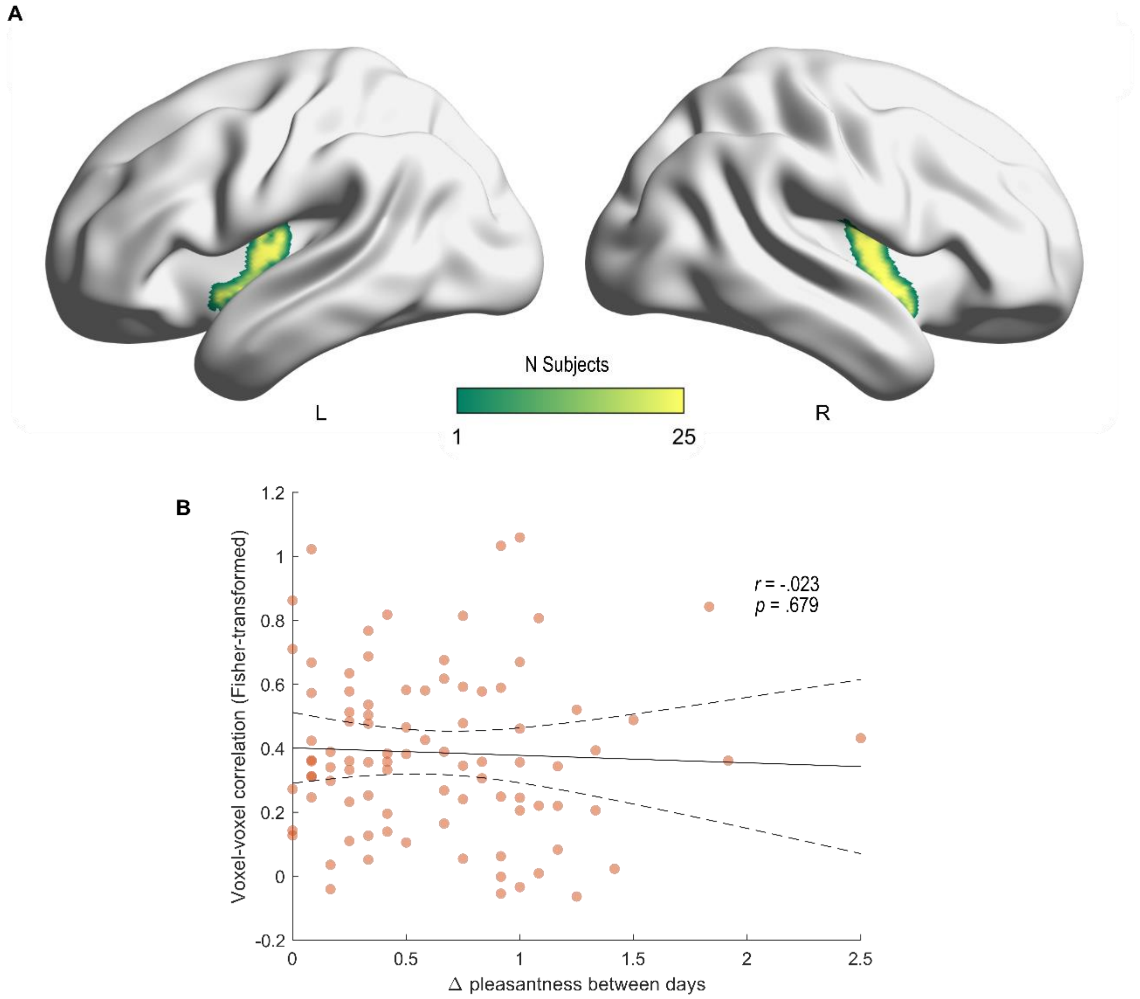


**Supplementary Figure 2.** Leave-one-subject-out ROI and pleasantness-induced change in flavour encoding patterns. **A)** Render of cortical surface and heatmap showing the number of subjects with specific sections of the insular cortex. Brighter colours mean more subjects had that part of the cortex. There is large variability in the leave-one-subject-out ROI, with some subjects extending further posterior and dorsally, with others more ventral and anterior. **B)** Plot of z-transformed voxel-voxel correlation against the change in rated pleasantness across days. There is no significant correlation between across-day correlation of activation patterns and change in rated pleasantness ($r$ = -.023, $p$ = .679). Dots indicate data points. Solid line indicates predicted value of regression model. Dotted lines indicate simultaneous 95% confidence bounds of the regression model.

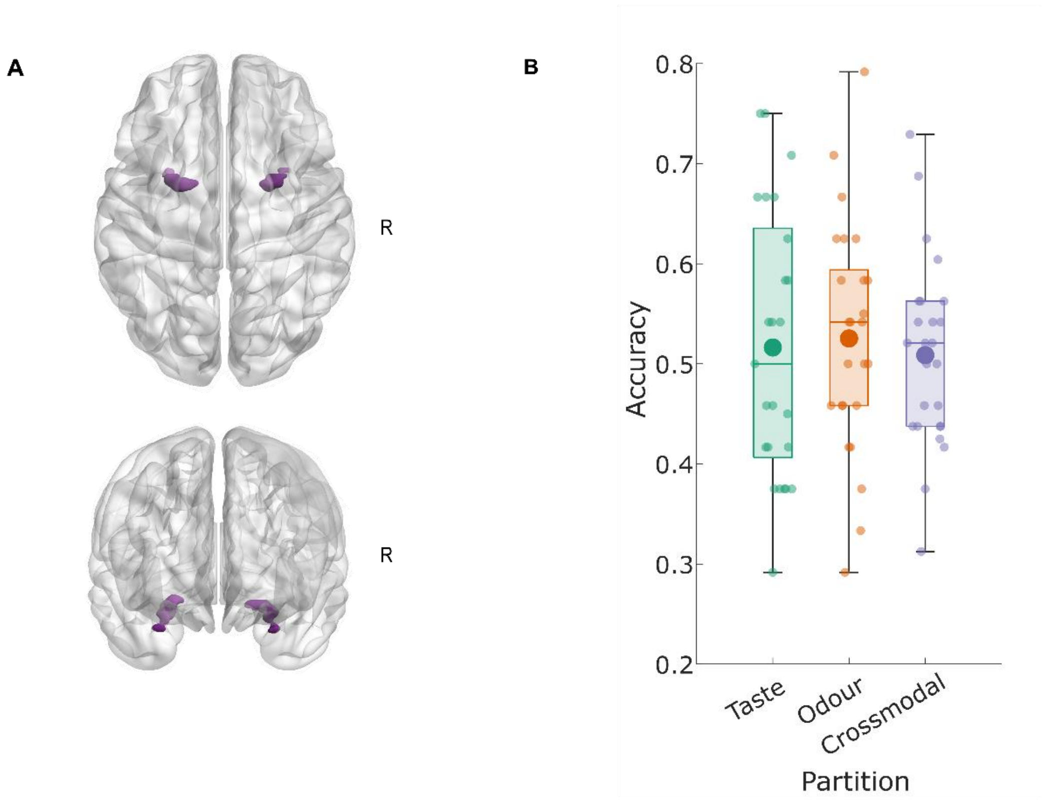


**Supplementary Figure 3.** ROI decoding in the piriform cortex. **A)** Axial (*top*) and coronal (*bottom*) views of the piriform ROI. **B)** Decoding accuracies in the piriform ROI. Decoding accuracy was not significantly above chance when distinguishing taste ($p$ = .2493) or odour ($p$ = .1384). Neither was crossmodal decoding significantly above chance ($p$ = .2808). Large dots signify means; small dots signify individual means; box limits indicate IQR; whiskers signify range as 1.5 times IQR; grey lines indicate subject-level changes across sessions.

**Table S1.**

Univariate unimodal BOLD activations against Artificial Saliva (ArtS)

| Peak Structure | Peak *x* | Peak *y* | Peak *z* | Volume ($mm^3$) | *Z* | $p_{FWE}$ |
|---|---|---|---|---|---|---|
| | | | **Odour > ArtS** | | | |
| **Piriform Cortex** | -18 | -2 | -14 | 414 | 5.56 | .0221 |
| **Piriform Cortex** | 26 | 0 | -16 | 258 | 4.17 | .0047 *SVC* |
| | | | **Taste > ArtS** | | | |
| **Mid-Insula** | -36 | -10 | 10 | 1019 | 5.16 | .0054 |
| **Mid-Insula** | 36 | -6 | 12 | 744 | 4.66 | .0126 |
| **Anterior Insula** | 40 | 4 | -8 | 234 | 4.46 | .0096 *SVC* |

SVC: small-volume correction on pre-defined coordinates (see **Methods**)

**Table S2.**

TFCE-corrected whole-brain searchlight crossmodal MVPA

| Peak Structure | Peak *x* | Peak *y* | Peak *z* | Volume ($mm^3$) | *Z* | $P_{TFCE}$ |
|---|---|---|---|---|---|---|
| **Mid Occipital Gyrus** | 38 | -74 | 32 | 14010 | 2.81 | .0024 |
| **Mid Frontal Gyrus** | 12 | 40 | -20 | 11617 | 2.03 | .0258 |
| **mOFC** | -18 | 28 | 16 | 335 | 1.78 | .0379 |
| **Cuneus** | 12 | -93 | 26 | 879 | 1.77 | .0382 |
| **Precuneus** | 4 | -60 | 32 | 748 | 1.77 | .0385 |
| **Inf Temporal Gyrus** | 50 | -62 | -12 | 1648 | 1.74 | .0401 |
| **Cuneus** | -6 | -82 | 20 | 1115 | 1.72 | .0431 |
| **mOFC** | -28 | 54 | -12 | 262 | 1.68 | .0467 |
